# Motor Planning Sensitivity to Affective Looming Sounds Within The Peri-personal Space: An Interplay of Exogenous and Endogenous Influences

**DOI:** 10.1101/2025.06.30.662313

**Authors:** Roberto Barumerli, Michele Geronazzo, Paola Cesari

## Abstract

Our brain maps the space immediately surrounding the body, the peripersonal space (PPS), to sharpen sensory-motor coordination whenever an object enters it. Within PPS, past research demonstrated how several factors influence motor readiness: from exogenous factors, such as body-object distance and stimulus semantics, to endogenous traits like personality traits. Nevertheless, most paradigms rely on vision or touch, relegating hearing to a supporting role and leaving auditory-only contributions unclear. Here, we tested whether affective content and individual traits modulate motor planning for looming sounds that stop within PPS. Thirty-three adults completed three auditory-only tasks in which positive, negative, or neutral sounds halted at five simulated distances from the participant’s ears (0.3–0.7 m). We recorded anticipatory postural adjustments, distance estimates, affective ratings, and sensory suggestibility via a questionnaire. Motor responses were largely anticipated as sounds stopped nearer the body, while delayed and less precise for semantic (positive or negative) than neutral sounds. Higher suggestibility predicted longer and more variable premotor latencies, particularly for non-semantic sounds. These findings show that auditory cues alone engage flexible sensorimotor mechanisms within PPS, where exogenous (distance, semantics) and endogenous (suggestibility) factors jointly shape motor readiness and spatial perception.

## 1. Introduction

To successfully navigate the environment, the brain maintains a dynamic representation of the body’s surrounding space by integrating sensory perception with action planning^1^. This representation, known as peripersonal space (PPS), enables rapid responses to potential threats, such as sidestepping an unseen car, and facilitates goal-directed behaviours, like catching a buzzing mosquito in the dark. Importantly, perception-action coupling within PPS boundaries is not fixed: they are shaped by both external environmental factors, such as the characteristics of stimuli entering the PPS (e.g., their nature, valence, or social relevance), and internally driven factors, including interoceptive accuracy and personality traits^2^. Prior literature offers a wide range of methodologies to investigate the factors influenced by the PPS, from reactive tasks that measure physiological responses probing underlying neural mechanisms to decision-making tasks that engage higher cognitive functions^3^. However, these exogenous and endogenous factors (i.e. environment-dependent configurations such as object distance and internal traits such as personality traits) and their effects on either low-level perceptual processes or high-level cognitive functions are often studied in isolation, limiting our understanding of how the brain operates as an integrated system^4^. Therefore, this study leverages auditory looming stimuli entering the PPS to gather evidence from physiological measurements, behavioural responses, and self-reported personality traits to examine how endogenous and exogenous factors modulate participants’ responses.

When an object enters the PPS, the brain must infer object characteristics from sensory inputs to enable a coherent interaction^2^. Much of our understanding of PPS comes from multisensory research, demonstrating that auditory, visual, and tactile cues converge to guide motor responses^3^. While hearing is often used as a facilitator and in support of touch and vision^5,6^, it is unclear how hearing alone provides a proxy for studying the perception-action integration of objects entering the PPS. Among these, looming sounds are particularly salient due to their ability to signal potential threats from a distance^7^. Empirical evidence shows that approaching sounds enhance spatial representation and trigger faster motor responses than those elicited by receding sounds^8,9^. Moreover, a sound’s meaning can influence the PPS’s size: negative sounds expand this boundary, so our defensive system is triggered by objects farther away than those perceived as neutral or positive sounds^10^. While these findings highlight the influence of both temporal, spatial features and semantic meaning on PPS, the role of individual variability in shaping auditory-motor coupling remains largely unexplored.

Anticipatory postural adjustments (APAs) offer a promising approach to investigate perception–action mechanisms, revealing how motor commands from the central nervous system prepare the body as a stimulus enters the peripersonal space^11^. APAs are early muscle activations that stabilise posture in preparation for movement. Their timing and contraction strength reflect the duration and facilitation that the central nervous system requires to process a perceptual event and initiate a motor response through the pre-motor cortex’s top-down modulation^12^. Unlike methods such as transcranial magnetic stimulation, hand-blink reflex elicitation, or electroencephalogram, APA measurement captures neural dynamics during natural movement, making it a more ecological and functionally relevant tool for studying sensorimotor integration in ecological environments^13^. In the auditory domain, APAs reveal feedforward motor responses: when looming sounds stop closer within the peripersonal space, APA onset occurs earlier, but sounds stopping outside PPS produce no such effect, demonstrating that this timing modulation is specific to stimuli entering the PPS^14^. Moreover, looming sounds with semantic content show valence-specific lateralisation: negative sounds strongly trigger APAs, reflecting a tight coupling between auditory perception and motor readiness, likely due to preferential engagement of the left motor cortex for unpleasant sounds and the right for pleasant ones^15–17^. However, existing studies have primarily focused on group-level effects, leaving open the question of how individual differences modulate APA timing within PPS boundaries.

Recent work investigating the link between perception and action shows its flexibility within PPS: sensory information is uncertain, therefore, people can actively interpret what they sense^18^. With such ambiguities, personal tendencies become evident: for instance, people with a high trait of anxiety lean more on their previous expectations when judging unclear motion, making their trait differences observable in perceptual decision making^19^. With regard to PPS flexibility, trait-level factors highlight this plasticity: higher trait anxiety enlarges PPS, as indicated by a stronger hand-blink reflex when the hand nears the face ^20^, whereas greater interoceptive accuracy narrows it, sharpening the boundary between the embodied self and the external world^4^. Although such studies relate PPS size to self-reported personality traits, they reveal little about how the perception-action coupling itself varies across individuals within PPS boundaries. Clues come from multisensory illusions within PPS, where highly suggestible people are more susceptible to the rubber-hand illusion^21^, possibly because enhanced multisensory integration and attentional engagement magnify their responsiveness^22^. When considering sensory suggestibility, defined as a person’s susceptibility to external sensory cues^23^, highly suggestible individuals might show differential activation within sensory motor pathways, especially auditory processing brain regions linked to spatial localisation and action preparation^24^. Together, these findings indicate that personal traits shape not only how far PPS extends but also how we prepare and act when stimuli enter this space.

In this study, we investigated how participant responses are modulated by endogenous and exogenous factors in response to looming auditory stimuli entering the PPS. We combined motor planning measures with an explicit distance estimation task. Participants were presented with three looming sounds, selected from the International Affective Digitized Sounds (IADS-2) database^25^, in addition to the pink noise stimulus, each stopping at five simulated distances within PPS. The affective properties of these sounds were validated using the Self-Assessment Manikin (SAM)^26^, while the Multidimensional Iowa Suggestibility Scale (MISS) quantified individual sensory suggestibility^23^. This dual approach allowed us to capture both rapid, pre-movement postural adjustments and conscious spatial judgments. To accommodate individual contributions of reaction times and perceptual estimates, we employed a Bayesian statistical framework, including beta regressions, thereby offering a flexible and robust analysis strategy that can be potentially extended to other dynamic perception-action paradigms^27^. By integrating neurophysiological measurements, crucial for understanding motor control in the premotor cortex, with explicit distance estimates, we aimed to elucidate how individual traits shape the interplay between sensory processing, decision making, and motor execution.

Therefore, we formulated two main hypotheses: (H1) when looming sounds entering the PPS carry semantic content (for example, a baby crying) versus no meaning (pink noise), experimental measures will show different levels of congruencies. In the cognitive distance estimation task, semantic sounds should be overestimated, whereas in the physiological task, the onset of initial muscle activation should be delayed, reflecting extra processing needed to decode meaning^28^. Second, we hypothesised that (H2) part of the inter-individual variability of the acquired physiological and cognitive data could be explained through the individual quantified level of sensory suggestibility, particularly when the incoming auditory stimuli carried distinct semantic connotations.

## 2. Results

With the general aim to investigate how auditory affective stimuli and individual suggestibility influence perceptual, spatial, and motor responses within the peripersonal space, we collected data from 33 participants (17 females; mean age = 23.3±3.2 years) who completed three auditory tasks and filled out a personality questionnaire. Participants self-reported normal hearing and no neurological or musculoskeletal impairments, all gave written informed consent, and the study was approved by the Ethics Committee of the University of Verona.

The tasks assessed distinct perceptual and behavioural responses to three auditory stimuli: two affective sounds selected from the IADS2 database^25^ (one with positive valence, ID 351 – Applause, and one with negative, ID 719 – Dentist Drill) and a neutral control stimulus (Pink Noise) without semantic content. An amplitude envelope following the inverse-square law was applied to simulate approaching sound sources^29,30^, starting 2.8 meters from the participant and moving at 0.7 m/s. Five stopping distances were defined relative to the participant’s ear, ranging from 0.7 to 0.3 meters in 0.1-meter steps (see Fig. 1a).

**Figure 1:**
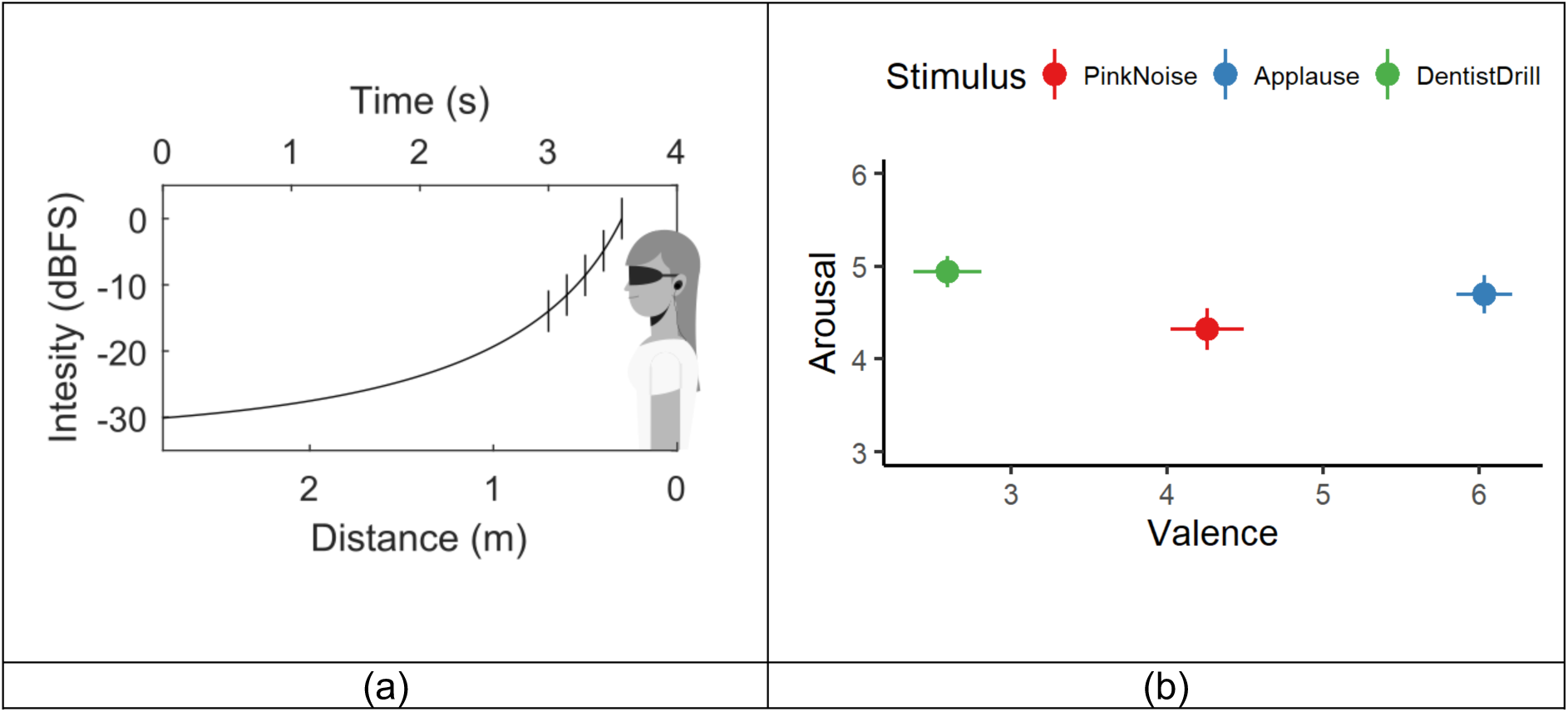
Characteristics of auditory stimuli and their affective evaluation. (a) Intensity curve of the experimental stimulus (vertical bars represent the five virtual stopping distances) along both time and distance axes. The participant’s ear canal was positioned at zero distance. (b) Scatterplot of mean arousal ratings as a function of mean valence ratings for three auditory stimuli. Error bars depict the standard error of the mean for both valence and arousal over participants.

Data were collected at the Biomechanics Laboratory of the University of Verona using motion capture, electromyography (EMG), and audio delivery systems. Kinematic data were recorded at 250 Hz using a VICON MX Ultranet system with reflective markers placed on the head, shoulders, and index fingers. EMG signals were recorded from the erector spinae muscles at 2000 Hz using a ZeroWire EMG system, synchronised with the motion capture via the Vicon control interface. Auditory stimuli were delivered through individually calibrated headphones, while the acoustic intensity was calibrated with a sound level meter. All data streams (including motion, EMG, and audio) were synchronised to ensure precise temporal alignment (see Methods for further details).

To combine neurophysiological data, behavioural measures, and self-reported suggestibility in a Bayesian model, we report results in three steps: (i) affective ratings of the sounds, (ii) estimates of each stimulus’s stopping distance, and (iii) reactive arm movements captured by EMG. The outcomes from distance estimation and muscle contraction timing are then related to individual scores of self-reported suggestibility obtained from the MISS questionnaire^23^.

### Affective evaluation of looming sounds

The affective evaluation was conducted using the SAM mannequin^26^, following the original methodology of the IADS2^25^, yielding strong concordance between our participants’ ratings and the original scores (see Fig.1b). An Aligned Rank Transform (ART) ANOVA^31^ revealed significant differences in valence across stimulus categories (F(2, 405) = 303.56, p < .001), but neither the distance factor (F(4, 405) = 1.99, p = .096) nor the interaction between stimulus category and distance showed significant effects (F(8, 405) = .61, p = .767). The partial eta-squared for the stimulus type was η_p_^2^=.6. Post-hoc contrasts computed by employing ART-C tests with Tukey correction^32^ indicated that valence ratings differed significantly among all three sounds, with negative stimuli rated lowest, neutral stimuli intermediate, and positive stimuli highest (see Fig.1b; all p < .001). Importantly, these differences in the valence score follow the intrinsic affective salience of the stimuli, as observed in the original work^25^. Similarly, statistical analysis of arousal values indicated significant main effects for both stimulus categories (F(2, 405) = 5.40, p = .005, η_p_^2^=.03) and stopping distance (F(4, 405) = 5.09, p < .001, η_p_^2^=.05), but not for their interaction (F(8, 405) = .63, p = .751). Post-hoc analysis revealed that the neutral stimulus was rated lower in arousal than the negative stimulus (p = .003), and the positive stimulus did not differ from both the neutral and negative stimuli (see Fig.1b; all p > .05). This supports the notion that perceived intensity modulates arousal and replicates findings that auditory intensity acts as a salient motion cue mediating the effects of looming sounds^8^. Further, all three sounds had tightly clustered arousal ratings (mean 4.65 ± 0.96), a much narrower spread than the full IADS arousal spectrum (mean 6.23 ± 2.16), ensuring comparable arousal across conditions.

### Stopping distance evaluation of looming sounds

The task required participants to estimate the distance between the body and the sound at its endpoint using their left arm as a scale as shown in Fig. 2a. Given the bounded support for responses that resulted in lack of normality, we ran a repeated-measures ART-ANOVA that revealed significant main effects of distance [F(4, 480) = 381.57, p < .001], stimulus type [F(2, 480) = 9.75, η_p_^2^ = .27, p < .001], and their interaction [F(8, 480) = 2.88, η_p_^2^ = .05, p = .004]. Post-hoc comparisons (ART-C tests with Tukey correction^32^) indicated that participants successfully discriminated ending distances across acoustic stopping points (mean ± se: .047 ± .004 m, p < .001 across distance levels). Further, neutral stimuli elicited significantly nearer estimates (.261 ±.008 m) than both positive (.314 ± .006 m) and negative sounds (.309 ± .014 m), with no significant difference between positive and negative stimuli (p > .05). Such results demonstrate systematic differences in responses when grouping them by semantic category suggesting an interaction between auditory object recognition and sound localisation^33^.

**Figure 2:**
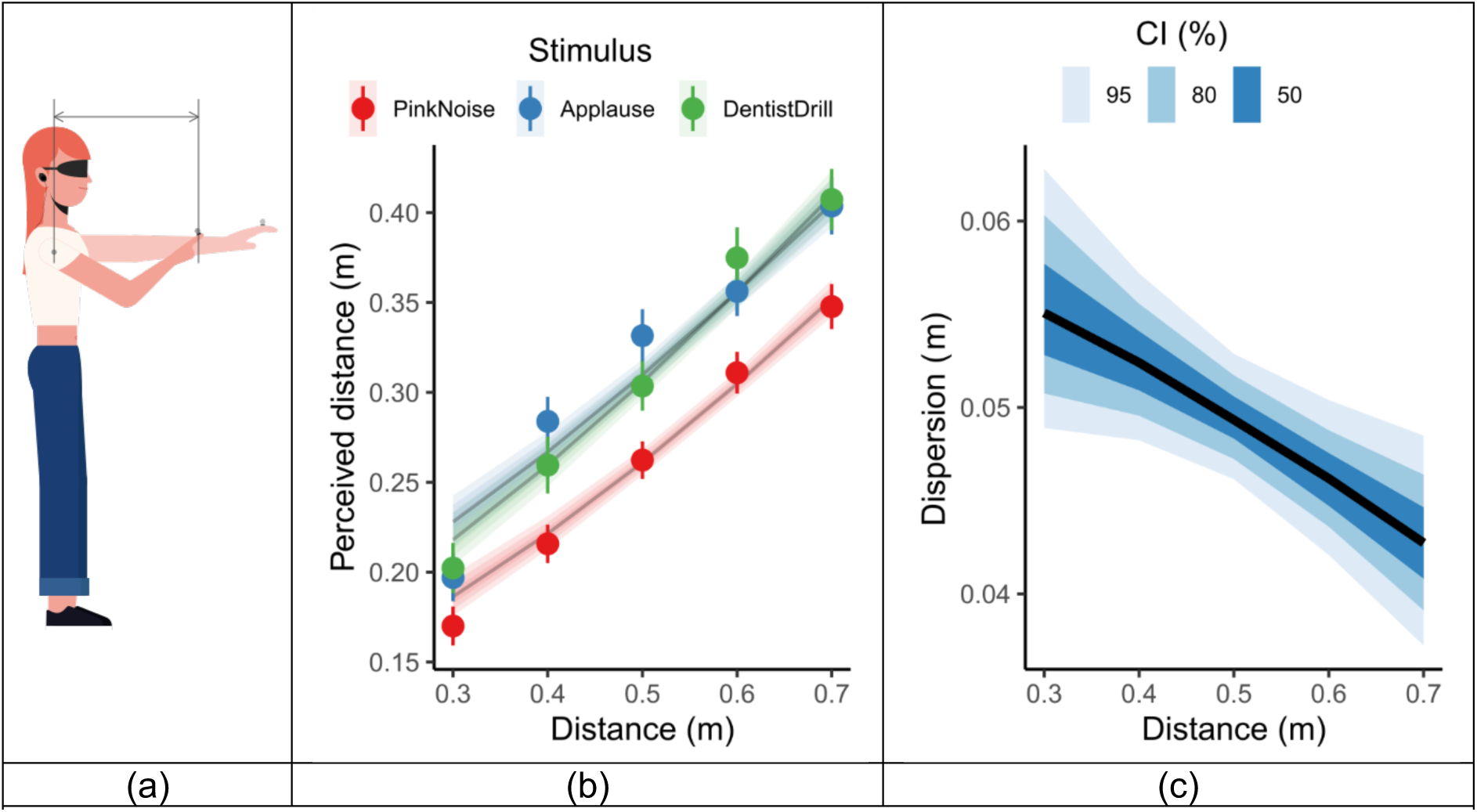
Distance estimation task and perceived distance of looming sounds. (a) Visualisation of a participant using her arm to provide the perceived final distance of the presented stimulus. (b) Estimated response of distance as a function of simulated ending distance for three auditory stimuli (Pink Noise, Applause, and Dentist Drill). Shaded areas represent 95% credible intervals of Bayesian beta regression while error bars report standard errors computed over participants. (c) Response dispersion as a function of distance, displayed with credible intervals at 50, 80 and 95% levels.

To further interpret these patterns in distance perception, we fitted a Bayesian beta regression model with weakly informative priors (see Sec.4.3), which supported the frequentist findings (see ribbons in Fig.2b). The posterior distribution for the slope of distance (.443 1/m, 95%-credible interval (CI) [.409, .478]) confirmed with strong evidence the participants’ ability to discriminate changes in auditory distance, with the 95%-CI not spanning zero. Additionally, systematic overestimations were observed for positive stimuli (.048 m, 95%-CI [.038, .058]) and negative stimuli (.045 m, 95%-CI [.034, .055]) compared to neutral stimuli. The Bayesian model allowed us to analyse response dispersion (i.e. empirical standard deviation visualised in Fig.2c) across experimental factors (distance and stimulus), demonstrating that dispersion linearly decreases with distance (-.032 1/m, 95%-CI [-.062, -.005]; p(v < 0) = .986), equating to a 1 cm decrease over the distance interval. This effect could stem from participants compressing their estimates near the shoulder because the arm’s finite length limits the measurement range.

### Premotor reaction time (pm-RT)

In this task, we asked participants to raise their arms as quickly as possible after the perceived end of the looming sound to measure postural adjustments as the contraction initiation of postural muscles^14^ (as shown in Fig.3a). We derived the premotor reaction time (pm-RT) as the activation time of the erector spinal muscles, and the analysis of variance (ANOVA) demonstrated significance for main effects of stimulus type (F(2, 480) = 90.754, p < .001) and distance (F(4, 480) = 381.571, p < .001), and their interaction (F(8, 480) = 2.881, p < .004). Similarly, for the perceived distance, pairwise t-tests with a Tukey correction revealed significantly faster pm-RT for neutral (176 ± .003s) against both positive (0.187 ± 0.002; p = .028) and negative sounds (0.191 ± .003; p < .001), highlighting the easier elaboration of such sounds (i.e., no need to identify the content); instead, the contrast between positive and negative remained inconclusive within the presented experimental data (p > .05).

We further investigated the trends using a Bayesian linear model (see Sec.4.3), which supported and extended the ANOVA findings (see Fig.3b). First, the estimated posterior distribution for the slope on the distance predictor was positive for neutral sounds (slope 0.046 1/s, 95%-CI [.031, .061]), indicating that greater distance is associated with increased pm-RT. Negative sounds systematically increased pm-RT timing compared to neutral sounds (intercept .010 s, 95%-CI [-.001, .022], p(v > 0) = .930), and positive sounds positively interacted with the distance term (.030 1/s, 95%-CI [.006, .054], p(v > 0) = .979). Second, we analysed how the variability of pm-RT, measured as standard deviation, changed across experimental factors (visualised in Fig.3c). For neutral sounds, distance introduced a negative effect (-.546s, 95%-CI [-.265, .164], p(v < 0) = .899), similar to the perceived distance, where uncertainty decreases for far distances. In contrast, semantic sounds showed increased uncertainty over distance, both for positive (1.201 1/s, 95%-CI [.188, 2.217], p(v > 0) = .975) and negative sounds (.983 1/s, 95%-CI [-.002, 1.961], p(v > 0) = .950).

### Suggestibility trait modules perception

Twenty-nine participants completed the Physiological Reactivity (PHR) subscale from the MISS questionnaire, a 13-item measure^23^. Unanticipated events during the experimental procedure led to incomplete data for four participants, which could not be recovered post-session. We used the PHR measure of self-assessed sensory suggestibility to evaluate the tendency to accept and act on perceived physiological states, with either reduced critical evaluation (low suggestibility) or independent judgment (high suggestibility). The subscale demonstrated good internal consistency (i.e. all items contributed coherently to the same construct), yielding Cronbach’s α = .810 with a 95 % confidence interval of [.581, .897]. Further, the distribution of raw scores was approximately normal (Shapiro-Wilk test, p = .654); therefore, we standardised the scores by subtracting the sample mean and dividing by the sample standard deviation.

We analysed how the PHR score might explain between-participants variability in both estimated distances and pm-RT. For both measurements, we included the normalised PHR score in the Bayesian models as an independent factor, as well as its interaction with remaining experimental factors (i.e., distance and stimulus semantics – see Methods).

For the perceived ending distance, the slope associated with the PHR score did not differ from zero (.004 m, 95%-CI [-0.021, 0.030], p(v > 0) = 0.625), and neither its effect on response dispersion (.001 1/m, 95%-CI[-0.006, 0.009], p(v > 0) = .628)). When evaluating how the suggestibility score influenced APA’s timing, we identified a relevant impact in predicting the accuracy and precision of pm-RTs as shown in Fig.4. In particular, the slope associated with the PHR score was positive (0.015 s, 95%-CI [0.003, 0.027]), indicating that individuals with higher suggestibility scores took longer to respond. The presence of semantics also impacted the response through the suggestibility score, with the positive semantics slowing down responses (0.0113, 95%-CI [0.006, 0.017]) as well as for negatives (0.0159, 95%-CI [0.011, 0.021]) in comparison with the sound without semantics (see Fig.4a). In addition, the effect of the PHR score on response dispersion (i.e., standard deviation) differed from zero only when considering the different types of stimuli (refer to Fig.4b): the variability increased only for the stimuli without semantics (slope = .004 1/s, 95%-CI [.001, .008], p(v > 0) = 0.987), while it remains constant for both negative and positive stimuli (*max*(*p*(*v*<*0*),*p*(*v*>*0*)) < 0.800). Interestingly, for participants with low suggestibility (PHR = −2), dispersion was significantly lower for non-semantic sounds than for semantic ones (p = 0.994 for v < 0). This indicates that these individuals responded more quickly and precisely when the sound lacked meaning. By contrast, for highly suggestible participants (PHR = +2), the dispersion did not differ across sound types (p = 0.832 for v > 0), showing that they remained slower and less precise regardless of the stimulus content.

**Figure 3:**
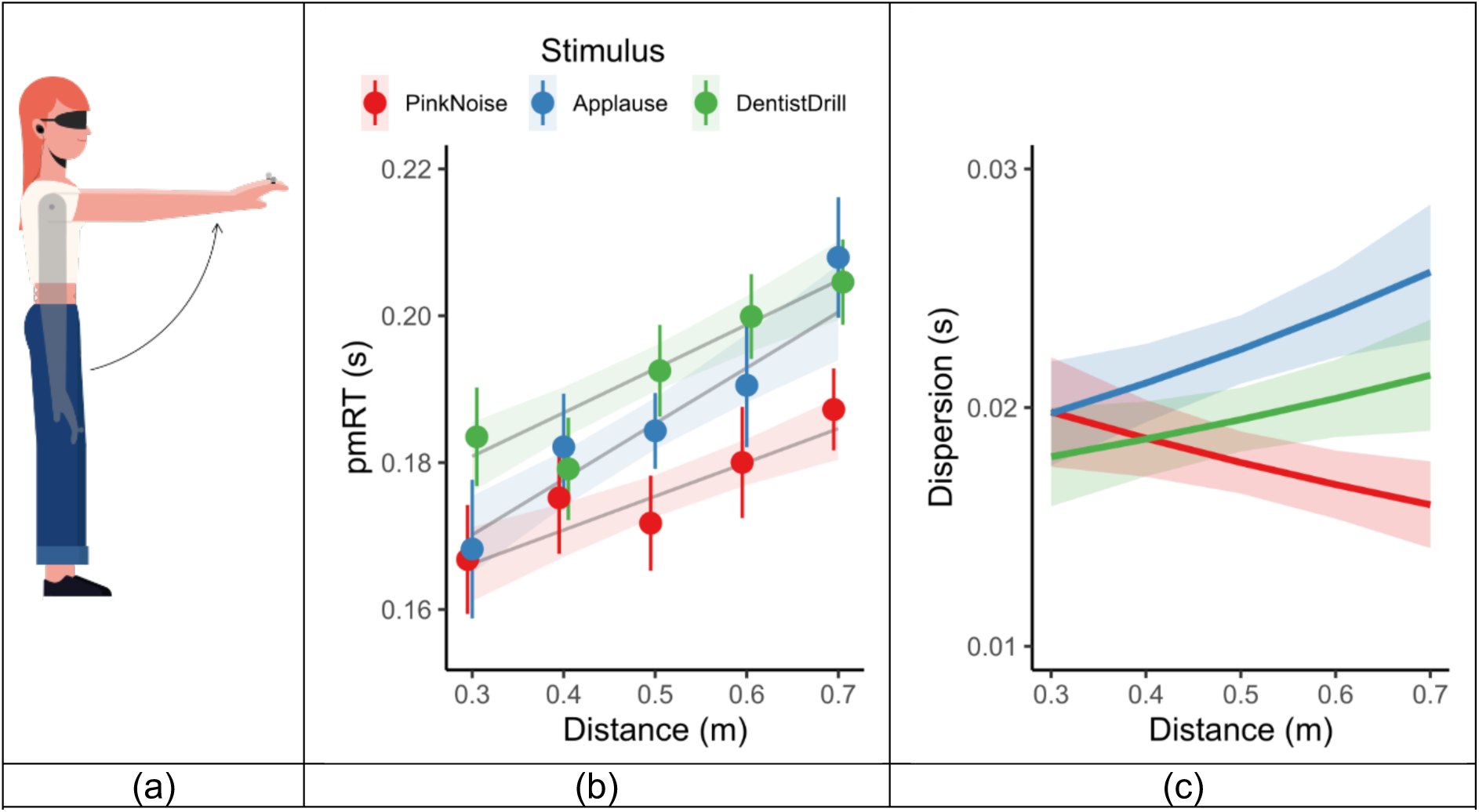
Premotor reaction time (pmRT) and its timing variability for looming sounds. (a) Visualisation of the participant raising her arm after the stimulus presentation. (b) Premotor reaction times as a function of simulated ending distance for three auditory stimuli (Pink Noise, Applause, and Dentist Drill). Shaded areas represent 95% credible intervals of Bayesian regression and error bars representing standard errors over participants. (c) Timing dispersion (i.e. standard deviation) as a function of distance, displayed with credible intervals at 95% levels.

**Figure 4:**
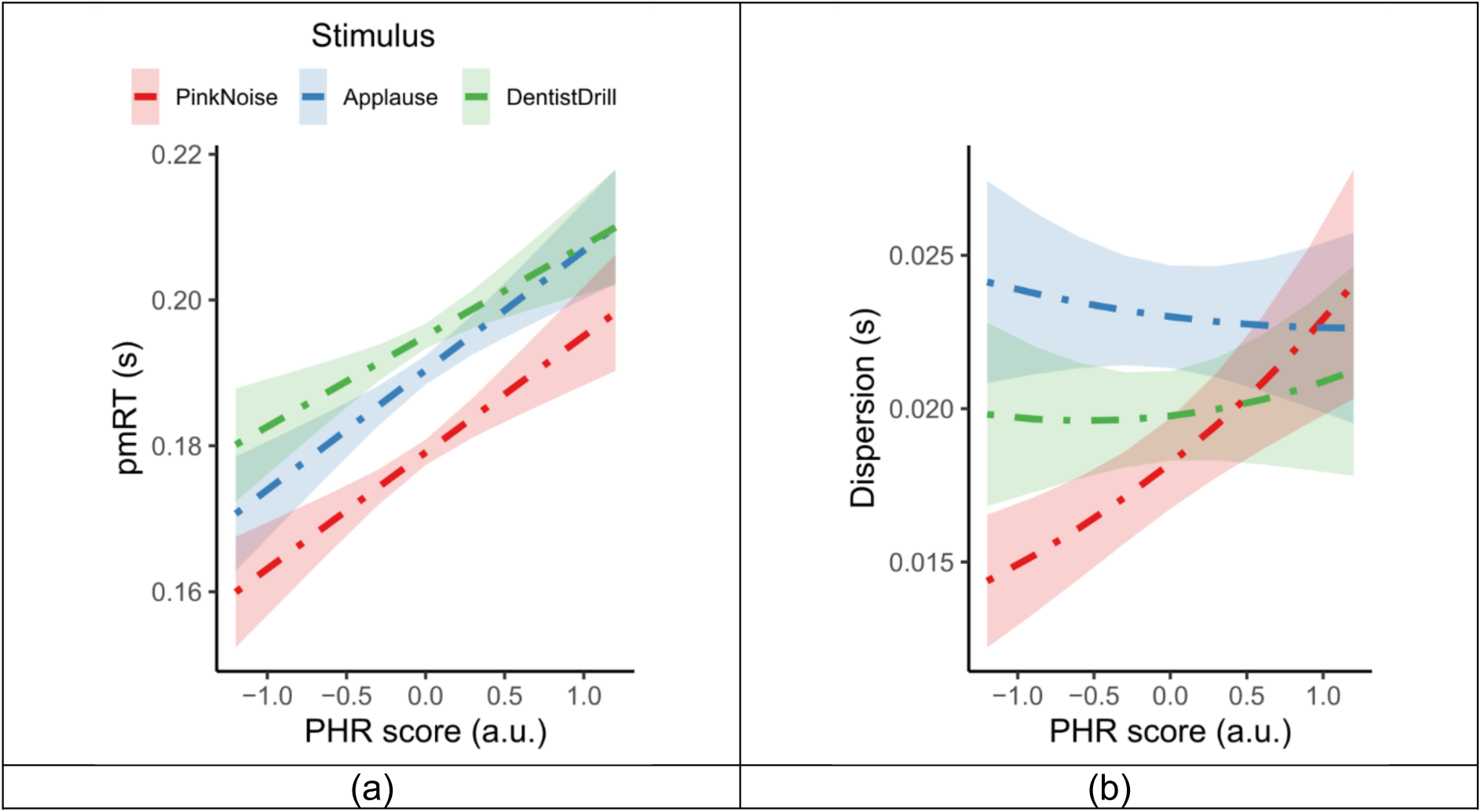
Influence of suggestibility on premotor reaction time and its dispersion. a) The plot shows the linear relationship between normalised PHR score and pm-RT response for three stimulus categories. b) The plot reports the effect of the normalised PHR score on response dispersion is shown separately for three categories. Lines represent linear regressions, and shaded areas indicate the standard deviation of slope estimates. PHR score is reported on a normalised scale (arbitray unit – a.u.).

## 3. Discussion

We jointly investigated endogenous and exogenous factors impacting proximity perception of looming sounds entering the PPS by combining physiological with behavioural measurements and self-assessed questionnaires within a simulated auditory environment. Overall, our results show that the endogenous factor (i.e. individual suggestibility) influences sensorimotor integration, linking this personality trait to the coupling between perception and action when a sound enters the peripersonal space. We confirmed the first hypothesis (H1): the estimation of looming sound-stopping distances (the cognitive task) and the timing of initial muscle activation in reaction to the looming sound-stopping distances (the physiological task) showed different levels of congruencies when comparing the exogenous factor associated to the presence of semantic (Applause and Dentist Drill) and without it (Pink Noise). For semantics sounds, we found an over-evaluation in the cognitive task and a retarded muscle activity in the physiological task. The second hypothesis (H2), instead, was only partially supported: individual levels of sensory suggestibility significantly influenced pm-RTs and their variability, while distance perception remained unaffected across all stimulus types. Interestingly, increased response variability in pm-RT measure was observed only for neutral (Pink Noise) stimuli, suggesting a defined interplay between semantic content and suggestibility in shaping reactive responses. Following, we discuss the results in more detail.

In this study, we relied on auditory distance perception to demonstrate that individualised looming sounds effectively modulate distance estimation and APA timing within the PPS, as confirmed by both behavioural and neurophysiological measures. In the distance estimation task, participants correctly estimated the stopping distances of the looming sound, demonstrating the participants’ ability to decode distance from rising intensity levels. Second, the measurement of pm-RT successfully replicated the trends observed in prior studies where sounds stopping nearer to the participant’s body elicit quicker reactions^15^. An important note is on the duration of our stimuli since it represented a confounder to sound intensity, however, the original study from Camponogara et al. 2015^14^ demonstrated that stimuli with a flat envelope and modulated duration did not elicit any modulation in the pm-RT indicating that participants did not perceived such control stimuli as approaching auditory objects. Further, the approaching stimuli were simulated through the inverse square law, following the concept that the intensity cue is the most important to elicit auditory distance^34^. Although the scientific literature indicates that additional cues, such as reverberation or spectral cues, should be considered to deliver an ecological percept^30^, the results obtained from both tasks demonstrate that participants were able to decode an estimate for auditory distance and use it effectively despite the simplifications introduced in stimulus rendering. This was evident both in the reactive task, where they modulated the pm-RT, and in the more cognitive task involving distance estimation.

Extending previous findings, our study examined how semantic content influences participant responses within the PPS, confirming anticipated relationships and uncovering novel effects. Participants accurately distinguished the valence of auditory stimuli, rating positive sounds higher than negative ones, with semantically neutral pink noise rated intermediately. Distance estimations and pm-RT revealed that meaningful (positive and negative) stimuli were perceived as more distant and elicited slower motor responses compared to pink noise. This may be due to additional cognitive processing required to decode semantic information, due to increased cognitive demand^35^, and with source localisation processing occurring more rapidly^28^ than identification^33^. While previous studies reported faster reactions to negative stimuli^10,36^, their use of tactile detection tasks and focus on PPS boundaries differ from our auditory-only, within-PPS design. The slower pm-RTs to both negative and positive stimuli may reflect inhibitory motor mechanisms associated with defensive processing of stimuli perceived as threatening due to their proximity, regardless of semantic content^8,17^. Similar effects have been observed in response to painful stimuli^37^ or unexpected, potentially threatening acoustic stimuli^38^. These results are consistent with evidence that premotor neurons support sensory-to-motor transformations^1,6^ and that threatening stimuli near the body activate cortical circuits involved in defensive behaviour, which can bias or suppress ongoing motor output to maintain a margin of safety^39^.

Prior literature has already linked how human defensive systems relate to personality traits, most notably anxiety and fear^40^. Therefore, we explored the possibility of explaining the variability observed in our data using the self-assessed MISS questionnaire, as responses might also be influenced by endogenous (internal) traits. Our results established suggestibility scores as a relevant predictor of participants’ timing in initiating APA (i.e. pm-RT). Specifically, participants with higher suggestibility showed slower reactions, consistent with a “freeze” response rather than a fight or flight reaction^41^. This pattern may reflect deeper engagement of the ventral auditory stream for decoding semantic content, which then modulates motor readiness in the dorsal pathway responsible for spatial and action processing^42^. Moreover, the presence of semantic content systematically increased pm-RT across participants, but notably, highly suggestible individuals (normalised PHR scores > 1) did not differentiate their pm-RT between negative and positive semantic sounds. Once again, such a result might indicate that increased suggestibility could affect top-down modulation from semantic interpretations, uniformly impacting pre-frontal cortex excitability and potentially reducing differentiation in motor responses based on sound valence^12,43^.

While the defensive-oriented role of PPS has been thoroughly explored, our results extend previous findings by demonstrating that response dispersion measures are also significantly modulated in the proposed within-PPS experimental design. Specifically, we observed that the presence of semantic content interacted with individual suggestibility, systematically influencing both response timing and precision. Recent studies have indicated that predictability and emotional context can alter the sharpness or variability of PPS boundaries^44,45^. In line with these observations, our data revealed that stimuli free of semantic meaning elicited increased variability in participants with higher suggestibility, suggesting broader or less precise PPS representations when meaningful contextual cues are absent. Conversely, when semantic context was present, response variability remained consistently higher irrespective of suggestibility, potentially due to additional cognitive processing demands related to decoding stimulus meaning and emotional significance^35^. This aligns with neural findings indicating high dorsal auditory stream engagement under conditions requiring explicit spatial evaluation without semantic clarity^28^. Importantly, this modulation of residual dispersion emerged exclusively in the pm-RT measure and not in the cognitively mediated estimated-distance task, likely because the slower, deliberate pointing response allows participants more cognitive control, reducing perceived threat and urgency^46^.

Despite the extensive study of PPS using visual and tactile inputs, the potential of auditory stimuli, especially in modulating motor readiness and spatial perception, remains an open and promising avenue for investigation. Our study highlights that auditory information alone can effectively shape PPS representations through loudness-based distance cues. Future research should further refine auditory simulations, incorporating realistic spectral and binaural cues beyond intensity changes to better approximate real-world acoustic dynamics^30,47^. Moreover, given the known interactions between interoceptive accuracy and PPS boundaries^4^, it would be valuable to integrate physiological measures (e.g., heartbeat tracking) into our paradigm to examine how individual differences in interoceptive sensitivity modulate auditory PPS processing and pm-RT. Additionally, our personalised auditory rendering approach lends itself naturally to implementations within immersive virtual reality^13^. Within these immersive scenarios, future studies could systematically investigate not only the multisensory integration of visual and auditory stimuli but also how imaginative suggestibility and sense of presence influence PPS boundary plasticity^48^. Exploring emotional processing further, dynamic measures of postural control^49^ alongside pm-RT and PPS measurements could clarify how affective auditory stimuli shape behaviours (e.g., freezing or avoidance) within PPS^16^. Lastly, future investigations through neuroimaging methods (EEG/MEG) could examine how auditory “what” and “where” auditory pathways^28^ interact to modulate sensorimotor integration in response to looming auditory stimuli^33^. Our Bayesian analyses and residual-based methodologies would provide effective tools for exploring how individual differences (e.g., personality, anxiety, interoceptive sensitivity) and stimulus properties jointly determine the variability in these neural and behavioural responses, offering deeper insights into the underlying mechanisms shaping PPS representations.

### Conclusions

Our findings demonstrate that auditory looming stimuli within the PPS effectively modulate postural adjustments and distance estimation, influenced by both exogenous (auditory distance and affective content) and endogenous (suggestibility) factors. We confirmed that stopping distances impacted our measures: as sounds stopped farther within peripersonal space, participants responded more quickly and estimated distance to be closer to the body. Instead, semantic information impacted experimental measures, slowing motor responses and increasing the perceived auditory stopping distance, likely reflecting additional neural processing demands to address the stimuli’s semantic content. Notably, individual suggestibility shaped both reaction timing and its dispersion, highlighting the importance of considering internal personality traits when studying sensorimotor integration. These findings reinforce the dynamic and personalised nature of PPS representations, extending our understanding of how the human brain integrates sensory, cognitive, and personality-driven processes to organise defensive and motor behaviours in response to approaching stimuli. In doing so, our study extensively explores the suggestibility role in sensorimotor integration with the aim of providing a blueprint for future research linking other personality factors, such as the Big Five personality traits^50^, to perception-action coupling within the PPS.

## 4. Methods

### Participants

Thirty-three right-handed adults (17 women; mean age: 23.4 ± 3.15 years) with self-reported normal hearing and no known neurological or musculoskeletal impairments volunteered for the study. Before participation, written informed consent was obtained from each individual. The study protocol was approved by the Ethics Committee of the Department of Neurosciences, Biomedicine, and Movement Sciences at the University of Verona, and all procedures were carried out in accordance with the Declaration of Helsinki.

### Stimulus generation

Three auditory stimuli were selected to represent distinct affective categories: Applause (positive valence), Dentist Drill (negative valence), and Pink Noise (neutral control). The emotional sounds—Applause (IADS ID 351) and Dentist Drill (IADS ID 719)—were sourced from the International Affective Digitised Sounds (IADS) database, which provides standardised ratings of valence and arousal^25^. The neutral stimulus, Pink Noise, was synthetically generated using MATLAB’s audio toolbox. Each stimulus was band-pass filtered (0.25–9.5 kHz) and RMS amplitude-normalised using the Pink Noise level as reference to ensure consistent loudness and spectral content across stimuli.

To simulate dynamic proximity, all sounds were transformed into looming stimuli through amplitude modulation by applying the inverse-square law of sound intensity decay, producing an exponentially rising intensity that mimicked a sound source approaching the body^29^. Specifically, the stimulus intensity ***I*** followed the physical relation 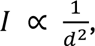 with ***d*** being the simulated distance between the source and the listener.

Each looming sound began at a simulated distance of 2.8 m, maintained a constant velocity of 0.7 m/s, and stopped at one of five predefined simulated endpoints within the auditory PPS (0.3, 0.4, 0.5, 0.6, and 0.7 m)^13,14^. The corresponding sound level increased from 65 dBA to 95 dBA across the trajectory^7,13^. A 15-ms raised cosine onset ramp and 20-ms offset ramp were applied to each stimulus to suppress acoustic startle responses and avoid off-response artefacts.

### Apparatus

The experimental procedures were conducted in the Biomechanics Laboratory of the Department of Neurological, Biomedical and Movement Sciences, University of Verona. Motion data were collected using a VICON MX Ultranet motion capture system (Oxford Metrics, UK), comprising eight Vicon MX13 cameras operating at a 250 Hz sampling rate. Reflective markers were placed on the head (midsagittal plane), shoulders, and index fingers to record postural and gestural kinematics. Custom MATLAB scripts enabled the automation of the experimental procedure and data collection.

Electromyographic (EMG) data were acquired from the erector spinae (ES) muscles using the ZeroWire EMG system (Aurion, Italy), with a sampling rate of 2000 Hz. The EMG multichannel analogue output was synchronised with the motion capture system via the Vicon MX control interface.

Auditory stimuli were delivered through Hefio One in-ear headphones, a research prototype that featured individual calibration of the ear canal for enhanced auditory rendering (refer to Geronazzo et al. 2023 for technical specifications^13^). An external audio interface (Saffire LE, Focusrite, UK) was used to amplify the audio signal. Sound intensity was calibrated using a CESVA SC-2c sound level meter, and playback was synchronised with kinematic data via the Vicon system. The Vicon MX control interface guaranteed synchronised recording of the audio signal with the motion capture and EMG data streams.

## 1. Procedure

### Task 1: Reactive Task

Participants were positioned at the centre of the recording space and blindfolded throughout the entire task execution to reduce visual interference in the responses. They sat upright with their hands resting comfortably along their sides. With this task, we measured APAs to quantify the timing of feedforward control in action anticipation, i.e., changes in muscle activation that occur before the initiation of a voluntary movement^51^. To elicit APAs, participants were instructed to quickly raise their arms forward in response to the cessation of each sound, triggering postural perturbations caused by dynamic intersegmental forces that shift the body’s center of mass forward, thus requiring preparatory muscle adjustments to maintain vertical posture. Auditory stimuli, designed to simulate looming sound sources, were presented binaurally and stopped at five simulated distances from the participant’s head. Each sound was presented with five repetitions per distance, resulting in a total of 75 randomised trials (3 sounds × 5 distances × 5 repetitions). Each block of 25 trials was followed by a 2-minute rest interval to minimise fatigue. Prior to the test, a training phase comprising 10 trials (5 distances as in the main tasks × 2 repetitions) with complex tones (100, 450, 1450, 2450 Hz) was conducted to familiarise participants with the response paradigm.

### Task 2: Auditory Distance Estimation

The second task was made to assess the spatial perception of auditory stimuli. The same three sound types and five distances were presented (4 repetitions per condition; 60 trials total) in randomised order. Still blindfolded, participants were instructed to extend their dominant upper arm horizontally as a proprioceptive metric for sound distance, with the shoulder being the most proximal point of reference and the middle fingertip as the most distal point of reference. Then, participants were instructed to approach their nondominant index finger towards the dominant upper arm until it physically reached the perceived stopping position. Experimental blocks, rest intervals, and training procedure were identical to Task 1.

### Task 3: Affective Evaluation of Sounds

The third task assessed the subjective emotional evaluation of the stimuli. The blindfold was removed, and participants used a touchscreen laptop with a screen of 15 inches to provide affective ratings using the Self-Assessment Manikin (SAM) scale^26^. Each 15 stimulus (3 sounds × 5 distances) was rated twice for two affective dimensions: Valence (1 = most negative; 9 = most positive) and Arousal (1 = least arousing; 9 = most arousing). Stimuli were presented in randomised order across all combinations of sound type and distance.

### Task 4: MISS

To gauge individual suggestibility to embodied auditory cues, participants were asked to respond to the Multidimensional Iowa Suggestibility Scale (MISS)^23^, a validated self-report inventory that assesses trait suggestibility across several domains rather than a single context (e.g., hypnosis). Among its five subscales, we focused on Physiological Reactivity (PHR), a 13-item measure of automatic bodily responses to internal or external cues (e.g., “Thinking about something scary can make my heart pound”), because it most closely aligns with our perceptual tasks.

## 2. Data Analysis and Preparation

### Task 1: Premotor Reaction Time

For each trial, the premotor reaction time (pm-RT) defined the initiation of muscle contraction (i.e. the timing of feedforward motor commands) by measuring the interval between the auditory stimulus offset and the onset of muscle activity in the erector spinae, recorded bilaterally^13–15^. The end of the auditory stimulus (trigger offset) was determined from the analogue trigger signal recorded during playback. After DC offset correction (mean of the first 200 samples), the signal was scanned to detect the global peak, followed by the signal descent below a dynamic threshold set at 10% of the peak amplitude. Instead, erector spinae onset was identified from EMG signals first low-pass filtered at 200 Hz (4th-order Butterworth) and then rectified. The envelope was computed via a secondary low-pass filter at 0.02 Hz (5th-order Butterworth). A dynamic threshold for activation was set as the mean baseline activity plus three standard deviations, calculated over a 100-sample window (i.e., 50 ms at 2000 Hz sampling rate) immediately preceding movement onset^14,51^.

### Task 2: Distance estimation

To estimate perceived sound location, we computed the Euclidean 3D distance between the participant’s pointing index finger and the ipsilateral shoulder, based on VICON motion capture data recorded during the final posture of the distance estimation task. The analysis window began 3 seconds after the sound offset, capturing the response phase. Within this window, a 15-sample moving average filter was applied to reduce measurement noise, and finger stabilisation was identified as periods where the finger’s 3D velocity remained below 0.01 m/s for at least 40 ms^13^. The final distance estimate was calculated as the Euclidean distance between the mean position of the stabilised pointing finger and the dominant-side shoulder.

### Task 3: Affective ratings

For each participant, the valence and arousal ratings collected by employing the SAM mannequin were averaged over repetitions, yielding one valence and one arousal score per stimulus type and distance pair.

### Task 4: PHR score

Participants’ raw PHR scores were obtained by summing the 13 items (each rated 1 to 5), which were used as a continuous predictor in the Bayesian models for Tasks 1 and 2.

## 3. Statistical analysis

The statistical inference workflow combined frequentist and Bayesian approaches to evaluate the effects of stimulus type and sound distance on behavioural and physiological responses. The section begins with an analysis of group-level differences using repeated-measures ANOVA. The analysis of differences is followed by Bayesian linear and generalised regression models tailored to the distributional characteristics of each dependent variable, allowing us to quantify uncertainty and test directed hypotheses on the relationships between measured quantities and experimental factors. Then, we explore how suggestibility traits, as captured by the MISS questionnaire, modulate perceptual outcomes.

### Analysis of the differences

Prior to statistical analysis, measurements were averaged over repetitions to guarantee stable estimates of individual performance and reduce the influence of trial-level noise or outliers. When reporting means and standard errors for each stimulus type, we computed them using a correction method for within-subjects^52^.

For the measurements of valence and arousal as well as for perceived distance metric, given the lack of normality on the residuals, we run a two-way aligned rank transform (ART) ANOVA^31^ on with the within-subject factors: stimulus type (Applause, Dentist Drill, Pink Noise) and distance (.3,.4,.5,.6,.7 m). Effect sizes (generalised eta squared, η²) were computed for all significant effects. For significant main effects or interactions, post hoc pairwise comparisons were conducted using sum-to-zero contrasts with Tukey correction for multiple comparisons. Post hoc contrasts were performed using the ART procedure for contrasts (ART-C)^32^. This method allows for valid nonparametric comparisons of factor levels following an ART ANOVA by applying linear contrasts to aligned-and-ranked data, while preserving the structure of factorial designs.

Statistical evaluation for the pm-RT metric was performed following a two-way ANOVA with two within-group factors, with five levels of distance and three levels for stimulus type (as for the previously described metrics). The data were processed and normalised per participant. Sphericity corrections (i.e., Greenhouse-Geisser adjustment) were introduced, and linear model residuals followed the normality assumption (p > .05). Effect sizes (generalised eta squared, η²) were computed for all significant effects. Post hoc comparisons were conducted using the estimated marginal means (EMMs) framework to follow up significant main effects and interactions identified in the ANOVA. Pairwise contrasts were applied to the EMMs for both stimulus type and distance, with adjustments for multiple comparisons using the Tukey method. Only contrasts with statistically significant adjusted p-values (p ≤ 0.05) were considered in the interpretation of results.

### Bayesian Statistical Analysis

With the aim of quantifying the relationships between experimental factors and measurements, we adopted a fully probabilistic modelling approach for inference based on Bayesian statistics^27^. For this analysis, we focused on the relationships between perceived distance, pm-RT, sound type, and the PHR score.

#### Model design

Bayesian models were fitted for each behavioural metric, mirroring the full factorial design used in the ANOVA. This allowed for direct comparability between frequentist and Bayesian approaches and ensured that all models captured the key experimental manipulations: stimulus type, distance, and their interaction, along with random intercepts for participant ID to account for repeated measures.

The analysis of perceived distance employed a Bayesian beta regression, which appropriately accounted for the bounded and non-Gaussian nature of this measure^53^. Perceived distances were expressed by pointing on the dominant arm, resulting in values between 0 and 1 m. Modelling both the mean and precision as a function of distance, stimulus type, and their interaction enabled the assessment of how both perceived proximity and response variability changed across conditions.

Instead, a Bayesian linear regression with a Gaussian likelihood described the pm-RT variability across experimental conditions. As with the distance model, this model included the full factorial fixed effects and participant-level random intercepts and modelled the residual standard deviation as a function of the same predictors to account for possible heteroscedasticity.

As a last step, we incorporated the normalised PHR score alongside the existing fixed effects (stimulus type, distance, and their interaction), as well as all two- and three-way interactions involving PHR. This extended structure was applied to both the mean μ and precision *φ* (or residual *σ*) components of the models, allowing the assessment of how individual differences in suggestibility influenced not only response magnitude but also response variability. Using the Wilkison notation and defining *f*(·) and *g*(·) as linking functions (i.e. identity for the Gaussian family, or *logit* and *log* respectively for a Beta regression^53^), the models followed this structure:

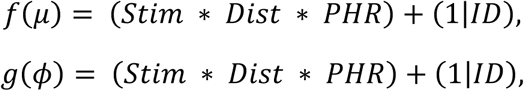

incorporating stimulus type (*Stim*), distance (*Dist*), and normalised PHR score (*PHR*) as fixed effects. Participant identifiers ***ID*** were included as a random intercept.

#### Parameter estimation and model evaluation

Model’s parameters were estimated using four Markov Chain Monte Carlo (MCMC) chains, with 3000 iterations per chain and 1000 warm-up steps. We increased iterations to 4000 and 2000 warm-up steps for models including PHR scores to reach convergence. Sampling employed Stan, a probabilistic programming language for Bayesian inference and statistical modelling^54^ and its default sampling scheme, No-U-Turn (NUTS), an adaptive form of Hamiltonian Monte Carlo designed to efficiently explore complex posterior distributions without manual tuning^55^. Default weakly informative priors (normal distribution, mean = 0, SD = 1). Convergence diagnostics (R^ < 1.01) and effective sample sizes were verified for all parameters^27^.

Model adequacy was evaluated using posterior predictive checks, drawing between 2000 samples from the fitted posterior distributions to compare model predictions with observed data across key experimental conditions. We compared the model’s average predictions and how much they varied from the real data. This approach enables the possibility to assess if the model reproduced the patterns observed in the data by visually inspecting summary plots and by employing aggregated statistics.

#### Probabilistic queries

Experimental effects were assessed through probabilistic queries of the posterior distributions of fixed effects. Directional and comparative hypotheses were tested with posterior probabilities for targeted hypotheses (e.g., whether the effect of stimulus Pink Noise was reliably smaller than Dentist Drill in proximity estimation). Posterior distributions were summarised using means and 95% credible intervals (95%-CI), giving a clear range of plausible effect values^27^. To further quantify certainty regarding the direction of these effects, we computed the probability of direction, representing the proportion of posterior samples consistent with the sign of the posterior mean. These metrics allowed us to interpret effect magnitudes and robustness without relying on binary significance thresholds.

To facilitate meaningful comparison between models using different likelihood families (i.e., Gaussian versus Beta regression), we standardised measures of variability. Specifically, for the beta regression, we computed the posterior standard deviation from the precision parameter to allow consistent interpretation of variability across different distributional families^53^.

We computed marginal effects to quantify how changes in predictors influenced outcome variables across their posterior distributions. For continuous predictors (e.g., distance), we estimated average marginal slopes, reflecting the instantaneous rate of change in the outcome per unit increase in the predictor. For categorical predictors (e.g., stimulus type), we computed average contrasts between categories, summarising differences in predicted outcomes across experimental conditions. Marginal effect estimates were summarised as posterior means and 95%-CI, providing robust population-level inferences integrated over observed covariate distributions.

Finally, we conducted post hoc exploratory analyses on model-derived parameters not directly estimated within the primary Bayesian models. Specifically, we extracted participant-level model estimates or their dispersion from posterior draws and then analysed their relationship with individual-level covariates (e.g., standardised PHR scores). For this procedure, we sampled 1000 draws and summarised them with mean and standard deviation over levels. These analyses allowed us to explore whether participant differences in suggestibility explained systematic variation in model uncertainty or sensitivity to experimental manipulations.

## 4. Software tools

Data acquisition and preprocessing were conducted using custom-written scripts in MATLAB. All subsequent statistical analyses were performed in R (version 4.4.2). For data manipulation and visualisation, we used packages including *data.table*, *ggplot2*, *car*, and *Rmisc*. Frequentist analyses (e.g., repeated-measures ANOVA, post hoc contrasts, and effect size estimation) were implemented using *afex*, *ARTool*, *emmeans*, and *effectsize*. Bayesian analyses relied on Stan via the interfaces provided by *brms* and *rstan*, complemented by *marginaleffects*, *tidybayes*, and posterior for marginal effects estimation, posterior exploration, model checking, and hypothesis evaluation.

## Funding declaration

This study was financially supported by the European Community Program NextGenerationEU in the form of a grant (PNRR M4-C2.1.1 PRIN 2022 project “S-TWIN” - 533 2022F9FWZ8 003 - CUP G53D23002840006) received by MG and PC.

## Acknowledgements

Figures 1a, 2a, and 3a include icons designed using resources from Flaticon.com.

## Author Contributions

**Roberto Barumerli:** Formal analysis; Visualisation; Writing – original draft; Writing – review & editing.

**Michele Geronazzo:** Funding acquisition; Conceptualisation; Methodology; Data Acquisition; Writing – review & editing.

**Paola Cesari:** Funding acquisition; Supervision; Conceptualisation; Methodology; Writing – review & editing.

All authors have read and approved the submitted version of the manuscript and agree to be personally accountable for their own contributions and for ensuring the integrity of any part of the work.

## Conflict of Interest

The authors declare no competing financial or non-financial interests.

## Additional Information

The authors declare that they have no competing financial or non-financial interests in relation to this work.

## Data availability

The datasets for the current study, as well as the analysis scripts, are available from the corresponding author on request.

## Notes

### Competing Interest Statement

The authors have declared no competing interest.

## References

1. Rizzolatti, G., Fadiga, L., Fogassi, L. & Gallese, V. The Space Around Us. Science 277, 190–191 (1997).

2. Bogdanova, O. V., Bogdanov, V. B., Dureux, A., Farnè, A. & Hadj-Bouziane, F. The Peripersonal Space in a social world. Cortex 142, 28–46 (2021).

3. Serino, A. Peripersonal space (PPS) as a multisensory interface between the individual and the environment, defining the space of the self. Neuroscience & Biobehavioral Reviews GG, 138–159 (2019).

4. Ardizzi, M. & Ferri, F. Interoceptive influences on peripersonal space boundary. Cognition 177, 79–86 (2018).

5. Canzoneri, E., Magosso, E. & Serino, A. Dynamic Sounds Capture the Boundaries of Peripersonal Space Representation in Humans. PLOS ONE 7, e44306 (2012).

6. Avenanti, A., Annela, L. C Serino, A. Suppression of premotor cortex disrupts motor coding of peripersonal space. NeuroImage 63, 281–288 (2012).

7. Neuhoff, J. G. Perceptual bias for rising tones. Nature 3G 5, 123–124 (1998).

8. Bach, D. R., Neuhoff, J. G., Perrig, W. & Seifritz, E. Looming sounds as warning signals: The function of motion cues. International Journal of Psychophysiology 74, 28– 33 (2009).

9. Ignatiadis, K. et al. Cortical signatures of auditory looming bias show cue-specific adaptation between newborns and young adults. Commun Psychol 2, 1–15 (2024).

10. Ferri, F., Tajadura-Jiménez, A., Väljamäe, A., Vastano, R. & Costantini, M. Emotion-inducing approaching sounds shape the boundaries of multisensory peripersonal space. Neuropsychologia 70, 468–475 (2015).

11. Bouisset, S. & Zattara, M. Biomechanical study of the programming of anticipatory postural adjustments associated with voluntary movement. Journal of Biomechanics 20, 735–742 (1987).

12. Maffei, G., Herreros, I., Sanchez-Fibla, M., Friston, K. J. & Verschure, P. F. M. J. The perceptual shaping of anticipatory actions. Proceedings of the Royal Society B: Biological Sciences 284, 20171780 (2017).

13. Geronazzo, M., Barumerli, R. & Cesari, P. Shaping the auditory peripersonal space with motor planning in immersive virtual reality. Virtual Reality (2023) doi:10.1007/s10055-023-00854-4.

14. Camponogara, I., Komeilipoor, N. & Cesari, P. When distance matters: Perceptual bias and behavioral response for approaching sounds in peripersonal and extrapersonal space. Neuroscience 304, 101–108 (2015).

15. Bahadori, M., Barumerli, R., Geronazzo, M. & Cesari, P. Action planning and affective states within the auditory peripersonal space in normal hearing and cochlear-implanted listeners. Neuropsychologia 155, 107790 (2021).

16. Bahadori, M. & Cesari, P. Affective sounds entering the peripersonal space influence the whole-body action preparation. Neuropsychologia 159, 107917 (2021).

17. Komeilipoor, N., Pizzolato, F., Daffertshofer, A. & Cesari, P. Excitability of motor cortices as a function of emotional sounds. PLoS One 8, e63060 (2013).

18. Noel, J.-P., Blanke, O. & Serino, A. From multisensory integration in peripersonal space to bodily self-consciousness: from statistical regularities to statistical inference. Annals of the New York Academy of Sciences 1426, 146–165 (2018).

19. Kraus, N., Niedeggen, M. & Hesselmann, G. Trait anxiety is linked to increased usage of priors in a perceptual decision making task. Cognition 206, 104474 (2021).

20. Sambo, C. F. & Iannetti, G. D. Better Safe Than Sorry? The Safety Margin Surrounding the Body Is Increased by Anxiety. (2013).

21. Marotta, A., Tinazzi, M., Cavedini, C., Zampini, M. & Fiorio, M. Individual Differences in the Rubber Hand Illusion Are Related to Sensory Suggestibility. PLoS ONE 11, e0168489 (2016).

22. Walsh, E. et al. Are You Suggesting That’s My Hand? The Relation Between Hypnotic Suggestibility and the Rubber Hand Illusion. Perception 44, 709–723 (2015).

23. Kotov, R. I., Bellman, S. B. & Watson, D. B. Multidimensional Iowa Suggestibility Scale (MISS) Brief Manual. (2004).

24. Looijestijn, J. et al. The auditory dorsal stream plays a crucial role in projecting hallucinated voices into external space. Schizophrenia Research 146, 314–319 (2013).

25. Bradley, M. M. & Lang, P. J. The International Affective Digitized Sounds (; IADS-2): Affective Ratings of Sounds and Instruction Manual. (2007).

26. Bradley, M. M. & Lang, P. J. Measuring emotion: The self-assessment manikin and the semantic differential. Journal of Behavior Therapy and Experimental Psychiatry 25, 49–59 (1994).

27. Gelman, A., et al. Bayesian data analysis third edition. Chapman and Hall/CRC (2013).

28. Ahveninen, J. et al. Task-modulated ‘what’ and ‘where’ pathways in human auditory cortex. PNAS 103, 14608–14613 (2006).

29. Neuhoff, J. G. An Adaptive Bias in the Perception of Looming Auditory Motion. Ecological Psychology 13, 87–110 (2001).

30. Zahorik, P., Brungart, D. S. & Bronkhorst, A. W. Auditory Distance Perception in Humans: A Summary of Past and Present Research. Acta Acustica united with Acustica G1, 409–420 (2005).

31. Wobbrock, J. O., Findlater, L., Gergle, D. & Higgins, J. J. The aligned rank transform for nonparametric factorial analyses using only anova procedures. in Proceedings of the SIGCHI Conference on Human Factors in Computing Systems 143–146 (ACM, Vancouver BC Canada, 2011). doi:10.1145/1978942.1978963.

32. Elkin, L. A., Kay, M., Higgins, J. J. & Wobbrock, J. O. An Aligned Rank Transform Procedure for Multifactor Contrast Tests. in The 34th Annual ACM Symposium on User Interface Software and Technology 754–768 (Association for Computing Machinery, New York, NY, USA, 2021). doi:10.1145/3472749.3474784.

33. Bizley, J. K. & Cohen, Y. E. The what, where and how of auditory-object perception. Nature Reviews Neuroscience 14, 693–707 (2013).

34. Middlebrooks, J. C. Sound localization. in Handbook of Clinical Neurology vol. 129 99–116 (Elsevier, 2015).

35. Beatty, G. F., Cranley, N. M., Carnaby, G. & Janelle, C. M. Emotions predictably modify response times in the initiation of human motor actions: A meta-analytic review. Emotion 16, 237–251 (2016).

36. Taffou, M. & Viaud-Delmon, I. Cynophobic Fear Adaptively Extends Peri-Personal Space. Front. Psychiatry 5, (2014).

37. Fossataro, C. et al. Anxiety-dependent modulation of motor responses to pain expectancy. Soc Cogn Affect Neurosci 13, 321–330 (2018).

38. Furubayashi, T. et al. The human hand motor area is transiently suppressed by an unexpected auditory stimulus. Clinical Neurophysiology 111, 178–183 (2000).

39. Graziano, M. S. A. & Cooke, D. F. Parieto-frontal interactions, personal space, and defensive behavior. Neuropsychologia 44, 2621–2635 (2006).

40. Perkins, A. M., Cooper, A., Abdelall, M., Smillie, L. D. & Corr, P. J. Personality and Defensive Reactions: Fear, Trait Anxiety, and Threat Magnification. Journal of Personality 78, 1071–1090 (2010).

41. De Vignemont, F. & Iannetti, G. D. How many peripersonal spaces? Neuropsychologia 70, 327–334 (2015).

42. Rauschecker, J. P. Where, When, and How: Are they all sensorimotor? Towards a unified view of the dorsal pathway in vision and audition. Cortex 98, 262–268 (2018).

43. Koban, L., Jepma, M., Geuter, S. & Wager, T. D. What’s in a word? How instructions, suggestions, and social information change pain and emotion. Neuroscience & Biobehavioral Reviews 81, 29–42 (2017).

44. Rossi Sebastiano, A., et al. Multisensory-driven facilitation within the peripersonal space is modulated by the expectations about stimulus location on the body. Sci Rep 12, 20061 (2022).

45. Matsuda, Y., Sugimoto, M., Inami, M. & Kitazaki, M. Peripersonal space in the front, rear, left and right directions for audio-tactile multisensory integration. Sci Rep 11, 11303 (2021).

46. Lu, J., Kemmerer, S. K., Riecke, L. & de Gelder, B. Early threat perception is independent of later cognitive and behavioral control. A virtual reality-EEG-ECG study. Cerebral Cortex 33, 8748–8758 (2023).

47. Baumgartner, R., et al. Asymmetries in behavioral and neural responses to spectral cues demonstrate the generality of auditory looming bias. Proc. Natl. Acad. Sci. U.S.A. 114, 9743–9748 (2017).

48. Jicol, C. et al. Imagine That! Imaginative Suggestibility Affects Presence in Virtual Reality. in Proceedings of the 2023 CHI Conference on Human Factors in Computing Systems 1–11 (ACM, Hamburg Germany, 2023). doi:10.1145/3544548.3581212.

49. Lebert, A., Chaby, L., Garnot, C. & Vergilino-Perez, D. The impact of emotional videos and emotional static faces on postural control through a personality trait approach. Exp Brain Res 238, 2877–2886 (2020).

50. John, O. P., Srivastava, S., & others. The Big-Five trait taxonomy: History, measurement, and theoretical perspectives. (1999).

51. Bertucco, M. & Cesari, P. Does movement planning follow Fitts’ law? Scaling anticipatory postural adjustments with movement speed and accuracy. Neuroscience 171, 205–213 (2010).

52. Morey, R. D. Confidence Intervals from Normalized Data: A correction to Cousineau (2005). TQMP 4, 61–64 (2008).

53. Cribari-Neto, F. & Zeileis, A. Beta Regression in *R*. J. Stat. Soft. 34, (2010).

54. Carpenter, B. et al. Stan: A Probabilistic Programming Language. Journal of Statistical Software 76, 1–32 (2017).

55. Hoffman, M. D. & Gelman, A. The No-U-Turn Sampler: Adaptively Setting Path Lengths in Hamiltonian Monte Carlo. Journal of Machine Learning Research 15, 1593– 1623 (2014).

